# Protein covariance networks reveal interactions important to the emergence of SARS coronaviruses as human pathogens

**DOI:** 10.1101/2020.06.05.136887

**Authors:** William P. Robins, John J. Mekalanos

**Affiliations:** Department of Microbiology, Harvard Medical School, 77 Avenue Louis Pasteur, Boston, MA 02115

## Abstract

SARS-CoV-2 is one of three recognized coronaviruses (CoVs) that have caused epidemics or pandemics in the 21^st^ century and that have likely emerged from animal reservoirs based on genomic similarities to bat and other animal viruses. Here we report the analysis of conserved interactions between amino acid residues in proteins encoded by SARS-CoV-related viruses. We identified pairs and networks of residue variants that exhibited statistically high frequencies of covariance with each other. While these interactions are likely key to both protein structure and other protein-protein interactions, we have also found that they can be used to provide a new computational approach (CoVariance-based Phylogeny Analysis) for understanding viral evolution and adaptation. Our data provide evidence that the evolutionary processes that converted a bat virus into human pathogen occurred through recombination with other viruses in combination with new adaptive mutations important for entry into human cells.

## Results and Discussion

The emergence of SARS-CoV and MERS-CoV as human pathogens is attributed to zoonotic infections in bats that transferred to civets and camels, respectively, while SARS-CoV-2 is similar to viruses isolated from both bats and pangolins implicating these species in the emergence of this historic virus (Al-Omari et al., 2019; Corman et al., 2014; Hu et al., 2017; Li, 2005; Wu et al., 2020; Zhang et al., 2020). The ∼30kb genome size of all SARS-related CoVs render sequence alignment and pairwise distance methods effective for phylogenic studies and predicting the likely source of such coronaviruses (CoVs). While nucleic acid sequence-based phylogenies are informative they clearly have limitations. Thus, we focused on variations in protein sequence to understand CoV evolution and the key functional interactions that drive adaption to new hosts or that influence transmission and pathogenicity.

We selected the conserved CoV proteins called 1a/1b, Spike (S protein), 3a, E, M, and N from a set of 850 viral genomes (Supplemental table S1). The alignment resulted in a 9639 amino acid consensus sequence with only 2% of sites being gaps with low coverage and 2% with low sequence conservation (Supplemental File F1). Because there are regions of diverged nucleotide identity in these viral genomes, our goal was to use amino acid identity to initially estimate phylogeny and then integrate that analysis with the identification of covariant residues within these six core viral proteins. Both SARS-CoV and SARS-CoV-2 are represented by a large collection of independent isolates from previous and current epidemics as well as from variants selected during passage in various laboratories (Supplemental Table S1) (Elbe and Buckland-Merrett, 2017). Similar to nucleic acid sequence-based analyses, our constructed phylogeny (Supplemental Figure S1) shows that SARS-CoV is closely related to CoVs found in civets and groups of bat CoVs that were previously suggested as likely ancestors (Li, 2005; Song et al., 2005). In contrast, SARS-CoV-2 is related to bat CoV RATG13 and also more similar to two bat CoVs, SZXC21 and SZC45, than other bat CoVs and its relationship to these in nucleotide similarity varies between large sections of the genome (Li, 2005; Paraskevis et al., 2020; Song et al., 2005). Five recently isolated pangolin CoVs identified as closely related to SARS-CoV-2 are also related to these three bat CoVs (Zhang et al., 2020) and our protein-based also analysis supports this observation. Thus, amino acid sequence-based analysis supports the conclusion that both bat and pangolin CoVs share or represent probable common ancestors of SARS-CoV-2 (Xia, 2020).

To survey the frequency of covariance among CoVs, we extracted pairs and groups (here designated ‘Clusters’) of covarying amino acid residues and organized these using a correlating tandem model that was set at different purity thresholds between 0.8 and 1.0. We selected a stringent purity threshold (0.96) to reduce noise based on our sampling size (Supplemental Table S2). We then applied force-directed mapping algorithms to visualize the relationships between these clusters of covarying residues and the CoV isolates from which they were derived (Jacomy et al., 2014). This graphing technique simulates repulsion between nodes (residues, clusters, and genomes) as well as attraction between edges (links between nodes) and then plots them to minimize both the complexity (fewer crossed edges) and to reduce edge lengths in two dimensions. Nodes are force-directed according to hierarchy and this orients these based on sequential linkage (edges) to clusters and then residues. Remarkably, we find that force-directed mapping of the linkage of covariance clusters readily organized all CoV isolates into groups that were consistent with phylogenic analysis (Figure 1 and Supplemental Figure S1). Thus, related genomes are forced into groupings in the middle of the graph surrounded by related clusters that are then flanked by networks of covarying residues. Furthermore, we provide every cluster and all residues identities by gene, genome, and host in a table and Gephi force map file (Supplemental Table S3 and Supplemental File S1) as well as an interactive website (https://sarscovresidues2.com/) We call this collective approach to understanding viral evolution ‘CoVariance based Phylogeny Analysis (CoVPA)’ in that it provides novel insights into the evolutionary origins of viral genomes.

**Figure 1.**
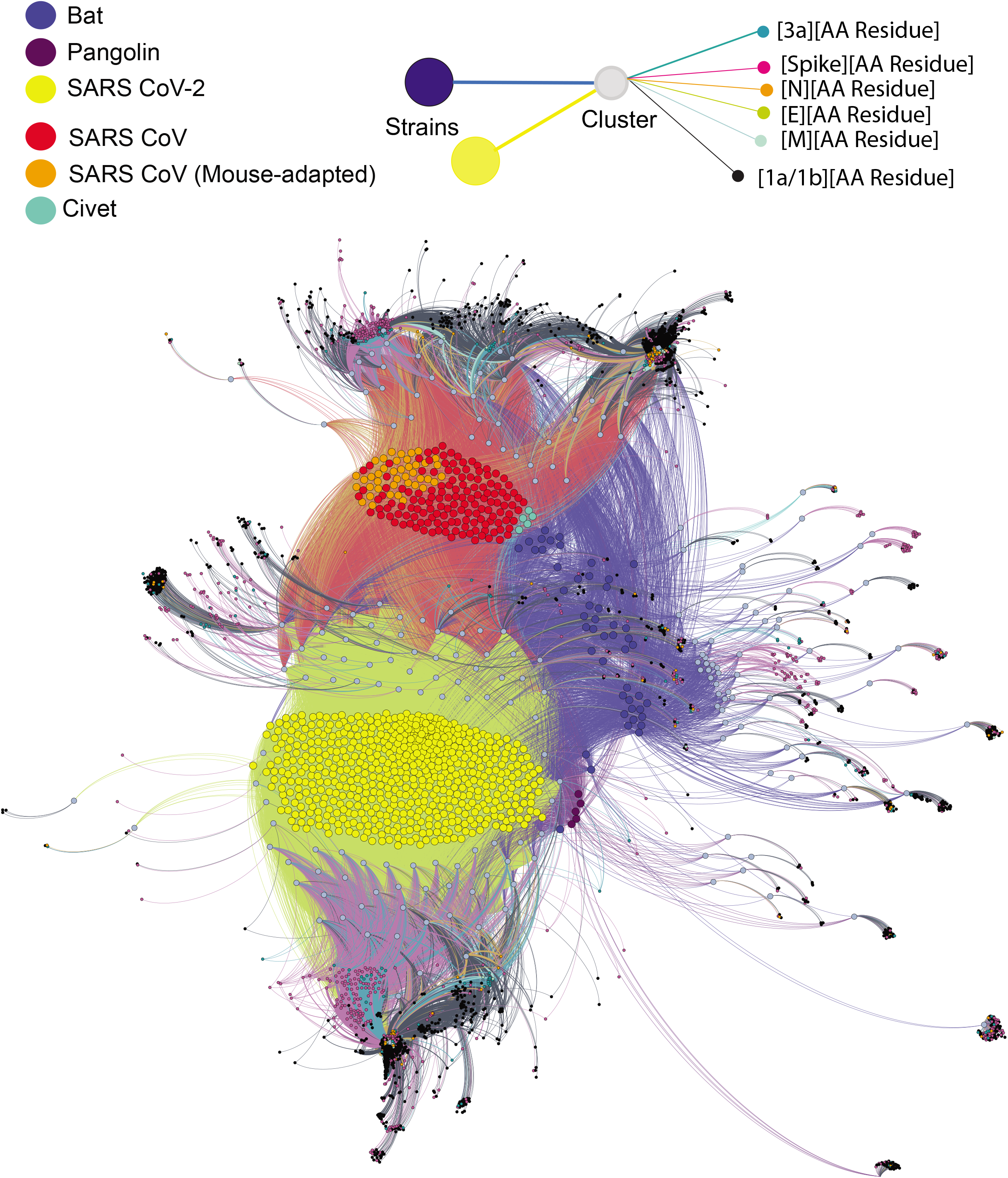
CoVPA map of covariant amino acid pairs and clusters of residues present in 850 SARS CoV genomes. Shown is a Gephi force map of the results of genome-wide covariance analysis which we have termed ‘<u>CoVariance based Phylogeny Analysis (CoVPA)</u>’. The data can be fully accessed via an interactive website (https://sarscovresidues2.com/) where a user can click on a map feature and site will then identify the protein residues, gene products, and viral genomes displaying the covariant residues or cluster selected. Complete tabular data sets of covariance data are also provided in Supplemental Table S3 and Supplemental File S1.

Some covariant clusters are ubiquitous to the entire grouping while others are exclusive to only one strain. We binned these clusters to compare covarying residues that are restricted to isolates from a given host species or a given group of CoVs. By comparing networks of clusters of covarying residues between distinct groupings, we could identify clusters that are restricted to various combinations of bat, civet, pangolin, and human CoV isolates. Thus, this annotated comprehensive dataset identifies all distinct residue identities that strongly covary in each CoV protein and the distribution of these in CoVs that include the human pathogens SARS-CoV and SARS-CoV-2. In its entirety, this provides a novel dataset that can be accessed through an interactive website (https://sarscovresidues2.com/) which allows one to explore the relatedness of CoV isolates based on conserved amino acid interactions that contribute to structure, protein-protein interactions, or biological function that multiple proteins drive interactively.

Both the civet and bat CoVs were previously identified as the closest relatives of SARS-CoV (Hu et al., 2017; Song et al., 2005) and these are positioned immediately adjacent to SARS-CoV in CoVPA mapping analysis. There are a few residues that are different between Civet CoVs (Cluster 26) and SARS-CoV (Cluster 65), but 103 residues are uniquely shared by both (Cluster 182). This supports previous findings that few genomic changes occurred between civet and the human-adapted SARS-CoV and that the majority of these polymorphisms likely arose during the initial transition from bats to civets (Song et al., 2005)). We conclude that these three clusters include residue identities that separate closely related bat CoVs from those enriched by passage and adaptation through other host species. The majority of these residues are in 1a/1b polyproteins, S, and 3a while covariance in E, M, and N is virtually absent.

Analogous to the civet and SARS-CoV relationship, three bat CoVs including RATG13 and five pangolin CoVs are all closely related to SARS-CoV-2 and are thus positioned in proximity in the CoVPA-network mapping approach because these share a significant number of covariant residues and clusters. In total, 10 clusters contain 1231 uniquely covarying residues specific to SARS-CoV-2 and the three most closely related bat CoVs. Covariant residues are highly represented in gene 1a encoding non-structural proteins (nsp1-nsp4), as well as the S protein and viroporin 3a (Supplemental Table S5). We surmise these enriched covariant residues include those that facilitated transmissibility into humans. Covariant residues are least represented in the RNA replication proteins, viroporin E, and membrane protein M. We did not find any cluster exclusive to all SARS-CoV-2 strains apart from those emerging in clinical isolates (Clusters 5 & 23) and instead find common overlapping clusters of covariant residues. For example, SARS-CoV-2 covariant residues are found to overlap with RATG13 (Cluster 183), or with RATG13 and pangolin CoVs (Cluster 194), or with the three bat viruses RATG13/COVCZ45/COVXC21 (Cluster 190), or with these three bat CoVs with pangolin CoVs (Cluster 197). As these CoVs share significant nucleotide similarity with another, this is not surprising; however, our covariance analysis also reveals key diverging features for the SARS-CoV-2 group that likely have biological significance. For example, there are covariant residues in the S protein that reside in its receptor binding domain (RBD), its amino-terminal domain (NTD), and in a structural region proximal to the furin cleavage site in its C-terminal domain (CTD). These SARS-CoV-2 covariant residues are likely key to the adaptation of this virus to new host species (see below).

We next focused on determining the covarying residues that allowed SARS-CoV-2 to emerge as a human pathogen by passage through intermediate hosts. To demonstrate the results of this analysis, we graphically show the stepwise progression and location of SARS-CoV-2 covariant clusters that overlap with those found in 48 bat CoVs and its most related bat isolate, RATG13 (Figure 2A, 2B, Supplemental Table S4). We plotted all covariant residue positions across multiple groups of CoV genomes to map their locations at the gene product level. The largest density of covarying residues in this progression to SARS-CoV-2 maps to both the NTD and RBD subdomains of S protein and the 3a viroporin protein. We conclude that enriched covariance networks between S and 3a allowed certain CoV isolates to adapt to growth in intermediary hosts and ultimately humans.

**Figure 2.**
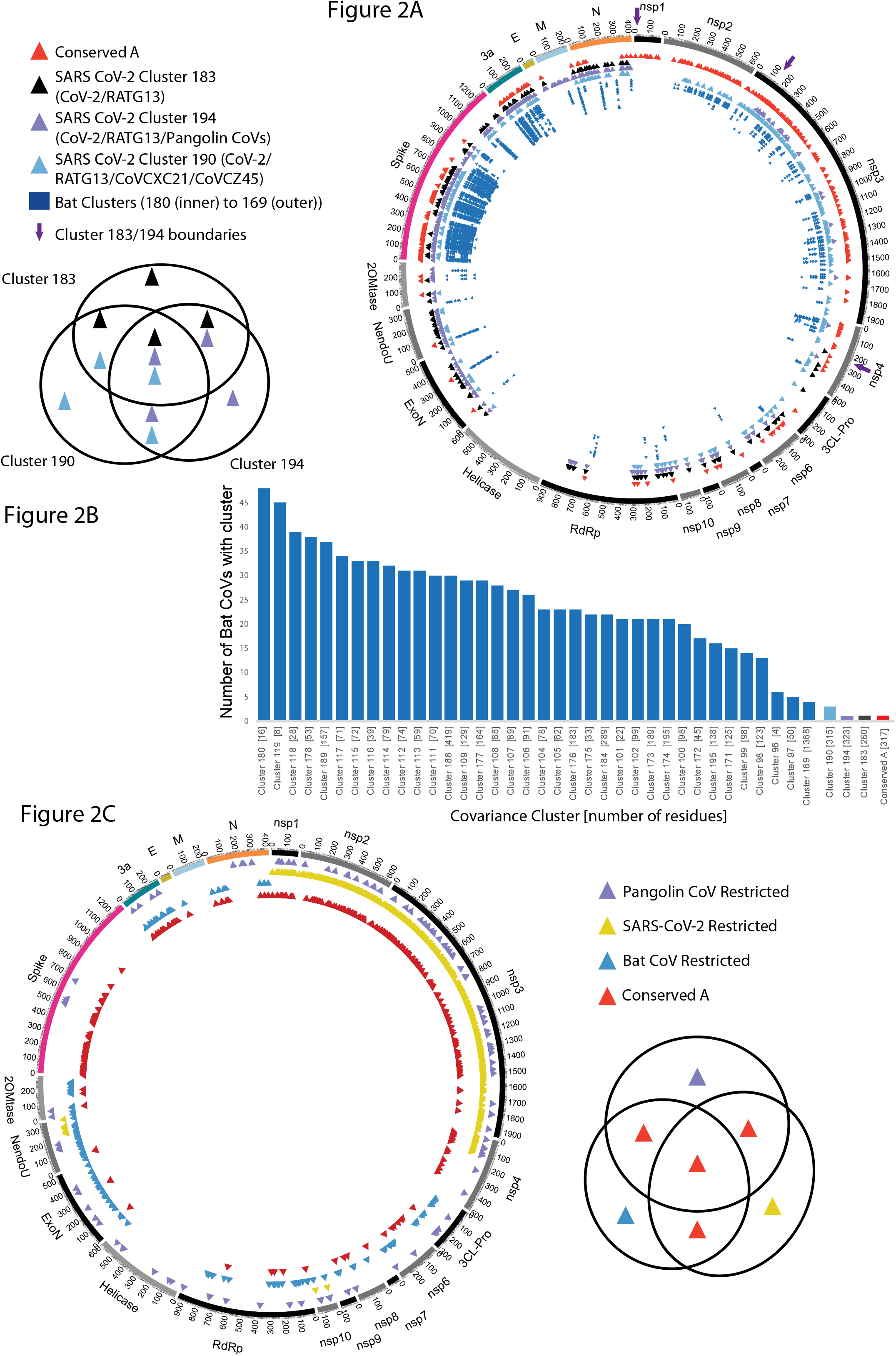
Plotting of predicted covariant residues in different CoVs by protein. **(A)** Mapped covariant residues in 1a/1b polyproteins, S, 3a, E, M, and N proteins from the bat CoVs found in SARS-CoV-2. Conserved residues found in other groups are mapped as a reference. **(B)** Graph showing a progression of Clusters and residue content more abundant in Bat CoVs to those more restricted to SARS-CoV-2 and closely related bat and pangolin viruses. **(C)** Residues found to covary that are restricted to categories of virus isolates. Notably, those restricted to SARS-CoV-2 are also shared with some Bat and Pangolin CoVs, but these residues are also uniquely exclusive to SARS-CoV-2 genomes. Those with overlaps and that are particularly well-conserved in between CoVs are shown as “Conserved A”; these covariant residues and clusters may have particular importance to the human infectivity of these pandemic viruses.

The role of covariant residues distinctly enriched in a specific host may be important in the context of adaption and infection. To separate such ‘restricted’ variants that are unique to CoVs from different species, we binned groups of covariants and then compared their mapped locations on CoV genomes (Figure 2B). Those classified as ‘restricted’ are found in clusters linked to specific groups and absent in other groups. Surprisingly both bat- and pangolin-restricted covariants are represented in distinct regions of the genome; for example, bat restricted residues are primarily located in nonstructural proteins encoded in 1b (2’O-Mtase, NendoU, ExoN) and enriched in 3a and N but not S or M. Clearly SARS-CoV-2 covariant residues belong to clusters linked to bat CoVs, but we found residues that are linked to this virus which are absent in other restricted categories (see below).

A large section in gene 1a (∼3000 AA, nsP1-nsP4) is unique to SARS-CoV-2 and distinct from the rest of the genome based on our covariance analysis. These contextual differences would not be readily apparent using standard nucleic acid-based single nucleotide polymorphism analyses. The identity and network of covariant residues in this region are unique based on matching mutations to those in bat viruses including those most similar such as RATG13 suggesting that this genomic region in SARS-CoV-2 experienced different selective pressures and evolutionary history (Figure 2A and 2B). The 3’-boundary of this region is aligned to a recombinational hotspot (Tagliamonte et al., 2020) and this suggests that hybridization of different CoV isolates likely led to the emergence of SARS-CoV-2. Though this region of SARS-CoV-2 is most similar to RATG13, the divergence of covariance residues between bat and pangolin CoVs here provides strong evidence that parts of SARS-CoV-2 emerged from an independent distinct source (Figure 2C).

In contrast to host-restricted covariant clusters, others are conserved covarying residues in other CoVs. For example, the Enriched Conserved A group contains residues that align in at least more than one independent category (Figure 2A and 2C); these enriched covariant groups include residues that are part of large networks of covarying residues (for example, in the nsp1-nsp4 region). This particular group was found to be also densely enriched in the NTD and RBD of the S protein, in much of 3a, and in both the CTD and NTD domains in N. The networks of covariant residues in the S protein may provide plasticity in protein domains important for receptor recognition and possibly escape from antibody responses given that the same regions are rich in dominant neutralizing epitopes (He et al., 2005; Qiu et al., 2005).

We examined the distribution and possible roles for the SARS-CoV-2 residues using the consensus sequence and by mapping these to domains in the S protein using the structural information present in the recently solved SARS-CoV-2 S trimer PDB file (Walls et al., 2020). In line with bat COVs, covariant residues in the S protein of SARS-CoV-2 are densely represented in both its NTD (residues 1-330) and its RBD (residues 333-527) (Supplemental Table S5). This includes residues between 438 and 506 that are part of the RBD interface with the ACE-2 receptor and also those between 333 and 526 that are thought to help stabilize the Beta-sheet and short connecting helices and loops that form the core of the RBD (Lan et al., 2020). Between 438 and 510 we identify 7 of the 16 residues shown to interact directly with ACE-2 (Lan et al., 2020) are covariant (Supplemental Figure S2). As covariance analysis detects residues that change with others, nearly all (6 of 7) are residues that differ between SARS-CoV and SARS-CoV-2. When aligned to known SARS-CoV RBD contacts with 80R, a potent neutralizing monoclonal antibody that has been co-crystalized with the S trimer (Hwang et al., 2006), we found that 13 of 19 residues contacted by 80R are covariants that differ between both SARS-CoV and SARS-CoV-2 (Lan et al., 2020). Notably, though much of S is highly conserved in structure and many CoVs share ACE-2 as a common receptor, it has been suggested that residues in the RBD-ACE-2 interface between SARS-CoV and SARS-CoV-2 are distinct enough to have arose by convergent evolution (Lan et al., 2020). Our analysis suggests that covariant residues likely influenced the receptor binding specificity of the S protein and perhaps also its ability to escape neutralizing antibodies.

To examine how the observed covariance compares to enriched mutations in the current epidemic, we examined 13,611 clinical strains sequenced since the emergence of the human epidemic from the GISAID database (Elbe and Buckland-Merrett, 2017). We identified covariant residue pairs most highly represented in clinical isolates and find these form small closed networks within and between proteins (Supplemental Figure 3A, Supplemental Table S6). These networks include some of the most abundant polymorphisms previously identified such 614 G/D in the S protein and 336 P/L in RdRp protein (Pachetti et al., 2020). Several residues located in the same small exposed flexible loop domain with 614 are predicted to be covariant in our 850 strains with residues in viroporin 3a as well as other residues in the S protein located between its RBD and furin cleavage site (Supplemental Figure 3B). The 3a and E viroporins of SARS-CoV are thought to be a K^+^ and Ca^++^ efflux channels and it is interesting that furin is proteolytically activated by these cations (Izidoro et al., 2010; Molloy et al., 1992). Furin cleavage of the S protein trimer is strongly implicated in its activation for membrane fusion events (Coutard et al., 2020). In this regard, we find residue 614 maps near the furin cleavage site in the S protein structure suggesting that 3a may be interacting with S during the process that leads to S cleavage and activation of membrane fusion. 3a and S protein have been found to form a complex but the nature of the interaction is still largely unknown at the structural level (Shen et al., 2005; Tan, 2005; Tan et al., 2004). Curiously residues in this exposed loop with 614 G/D are found to also covary with residues in the NTD of the S protein and in 3a suggesting there is an evolutionary relationship between 3a protein, S domains proximal to the furin cleavage site, and its NTD.

Our observed covariance in the 3a viroporin was mapped to predicted functional domains of the S protein (Supplemental S3B, S4A). The 3a protein of SARS-CoV is predicted to assemble as both a homodimer and a tetramer that form a membrane channel (a viroporin) in the cell and possibly the viral particle membrane (Ito et al., 2005; Lu et al., 2006). We noted covariance in distinct separate domains of 3a. There are covariant residues within a more variable NTD region (the first 30 residues of 3a) which is predicted to be surface-exposed as well as in other predicted extracellular and cytoplasmic loops (Supplemental Figure S4A). Antibodies to 3a and its N-terminal ectodomain from convalescent-phase SARS-CoV patients are commonly found (Qiu et al., 2005; Zhong, 2006) and the covariant residues that map here overlaps with a hypervariable region predicted to be more antigenic (Supplemental Figure S4B, S4C, S4D). We also identified a cluster of covariable residues in the CTD between 168 and 189; there is no role for this subdomain though it partially overlaps between residues 160 to 173 that contain putative intracellular protein sorting and trafficking motifs (Huang et al., 2006; Tan et al., 2004). In addition to a role in intracellular vesicle formation through interactions with Caveloin-1, the 3a viroporin of SARS-CoV protein is known to cause activation of the NLRP3 inflammasome, NF-kB activation, and chemokine induction as well as efficient viral release from infected cells (Castaño-Rodriguez et al., 2018; Chen et al., 2019; Freundt et al., 2010; Lu et al., 2006; Padhan et al., 2007). Thus identifying 3a as a hotspot for covariance and mutational selection may have relevance to the severe inflammation seen in COVID-19 disease and suggests that the 3a protein is a potential new target to consider for SARS-CoV-2 immunoprophylaxis.

The mutational history of genes are seldom random throughout evolution and display biases in regard to purine or pyrimidine substitutions that may reflect selective pressures such as polymerase fidelity, codon usage, and functionality of encoded gene products (Reviewed by Lyons and Lauring, 2017). We wondered whether these biases might provide further evidence of the selective pressures experienced by covariant residues during the evolutionary histories of SARS CoV virus families. We measured the frequency of transition (purine-purine, pyrimidine-pyrimidine) and transversion (purine-pyrimidine) signal nucleotide polymorphisms (SNPs) between SARS-CoV-2, RATG13, and one of the closely related Pangolin genomes analyzed in this work, PanCoV-GX-PL4, and found similar to other studies (Lam et al., 2020) a bias against transversions. We did not compare these with Guangdong Pangolin CoVs GD/P1L and GDP2 from Lam et al, 2020 that possesses a RBD even more similar to SARS-CoV-2 due to an incomplete sequence at the time of this work. The frequency of this bias (ratio of transition to transversion mutations) between the genomes of SARS-CoV-2 and RATG13 was approximately 2.0; in contrast, when comparing these two CoVs to PanCoV-GX-PL4 this bias was 7.5 (Figure 3). We interpret this result as evidence for the very recent evolutionary divergence between SARS-CoV-2 and RATG13 while PanCoV-GX-PL4 is clearly more evolutionarily distant relative to RATG13 and SARS-CoV-2.

**Figure 3.**
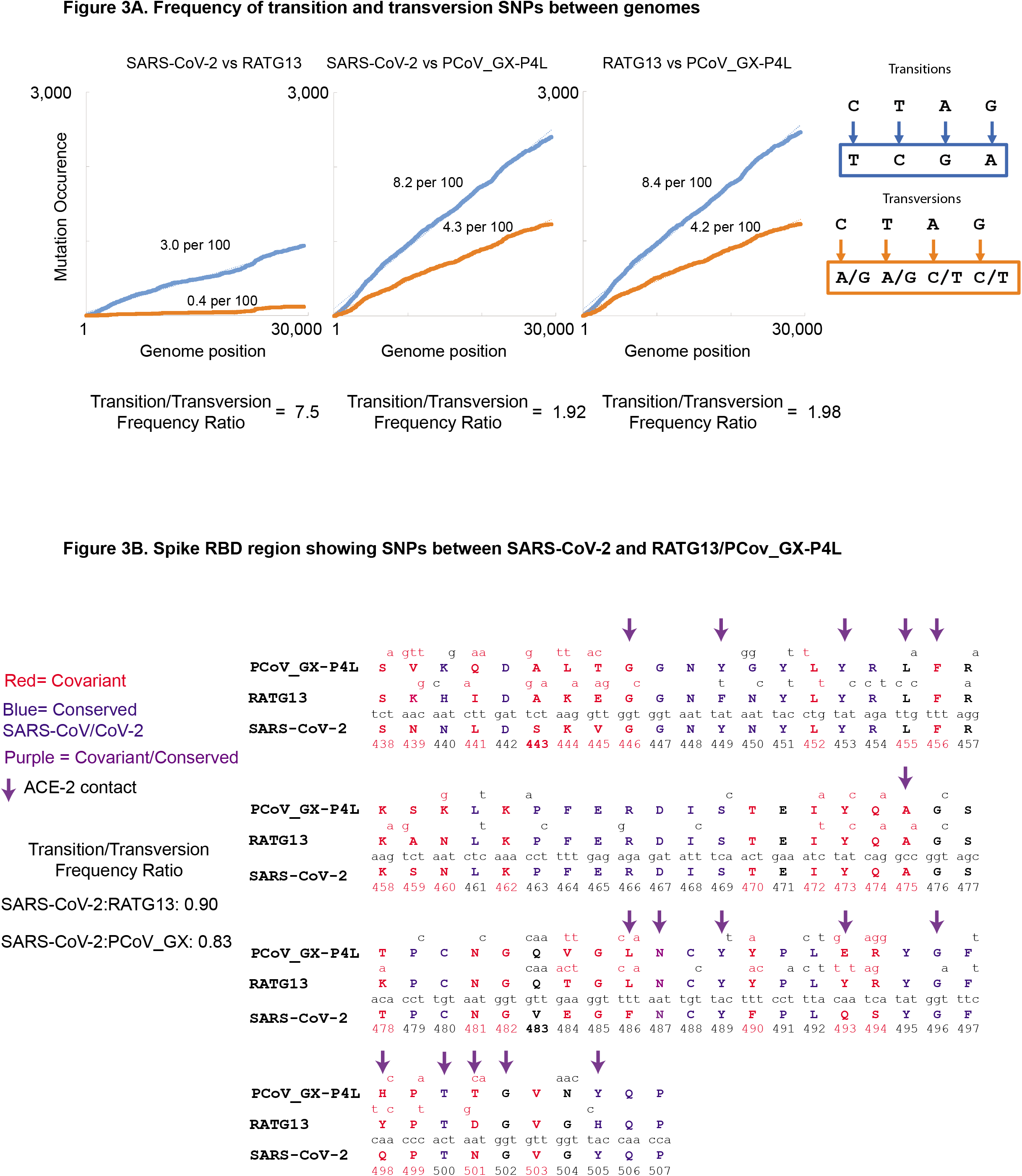
Measurement of the frequency of transition and transversion SNPs between SARS-CoV-2, RATG13, and PCoV-GX-P4L. **(A)** Plots separately showing the frequency of transitions and transversions in SARS-CoV-2, RATG13, and PCoV-GX-P4L. **(B)** The alignment of both amino acid and nucleotide identities RATG13 and PCoV-GX-P4L to SARS-CoV-2 within the RBD domain of S protein. Covariant residues are colored red and conserved residues blue. Arrows indicate determined contacts between ACE-2 and RBD residues of the S protein.

Though the rate of SNP accumulation across the 30Kb genomes appears at a gross level to be linear, there are regions in genes where SNPs are densely represented relative to other regions. In the RBD of the S protein, a recognized region enriched for covariant residues in this work, we find the frequency of transversion SNPs exceeds that of transition SNPs (Figure 3b). This 8.3-fold change in the ratio of transition to transversion, 7.5-fold to 0.90-fold, within the region encoding the RBD involves many codons encoding covariant residues. For example, 12 of 16 residues that changed in this region are in covariant residues in SARS-CoV-2 and many of these have accumulated 2 to 3 SNPs. Furthermore, non-covariant residue 483, a codon entirely absent in SARS-CoV, is found to differ at all 3 codon positions as transversion SNPs when SARS-CoV-2 is compared to RATG13 and PanCoV-GX-PL.

The differences in the RBD of the S proteins of SARS-CoV-2 compared to RATG13, and pangolin viruses such as PCoV_GX-P4L has been previously attributed to recombination events that occurred between related ancestral viruses (Huang et al., 2020; Li et al., 2020; Zhang et al., 2020). However, we show here that the RBD of SARS-CoV and SARS-CoV-2 possess a pattern of conserved covariant residues (Supplemental Figure S2), that were likely evolved in both these viruses under selection pressures that cannot be easily attributed to recombination alone. The inherent diversity of covariant residues identified in this analysis is demonstrated by the observation that only one covariant residue (N487 of the S protein) is also conserved in identity between SARS-CoV and SAR-CoV-2 (Supplemental Figure S2). Of the 16 residues in SARS-CoV shown to interact with the human ACE-2 receptor (Lan et al., 2020), 7 are covariant residues and an additional 5 are adjacent to covariant positions. These residues are predicted to covary with others in the RBD and residues primarily within S and 3a in both SARS-CoV and SARS-CoV-2. We hypothesize that the purifying selection imposed by these and other covariant networks resulted in the enhanced selection of radical changes in the covariant residues in the RBD of SARS CoV-2 that we observed. Thus, the observed co-variability of residues in the NTD and RBD of S as well as within the 3a protein is more easily explained by selective processes that can drive mutations in multiple sites within the same or different proteins rather than simple recombination events that would produce dramatic but uncoordinated changes in the structure of proteins.

Though CoVPA presents covariance networks with numerous residues in SARS-CoV-2, there are a few small clusters identified in this work that likely reveal host-adaptation of CoVs. For example, Cluster 62 identifies 5 residues that emerged in SARS-CoV during its further passage and adaptation in the mouse model. In the context of human CoV diseases, we also identified one covariance network of eight residues in the S protein (Cluster 126) found only in SARS-CoV and SARS SARS-CoV-2 and the most closely related civet, pangolin, and bat CoVs (Supplemental Figure S5A, S5B). Though some of these residues are missing from the available models of the S trimers from both SARS-CoV and SARS-CoV-2, we find residues Q23, H66, and V213 are proximal to one another when comparing the tertiary NTD structures of both (Supplemental Figure S5B, S5C). The relative position of two other covariant residues L54 and M153 in the NTD is difficult to determine in that they reside within disordered and exposed regions in the cryo-EM S trimer reconstructions of SARS-CoV and SARS-CoV-2. There are no known interactions or roles for these protruding NTD regions of the S trimers. Because the NTD of the S protein is especially enriched in covariant residues, we propose that host- or immune-driven selective pressure has occurred within this subdomain. In combination, covariant residues in S protein appear to have emerged independently and are enriched in viruses that caused two independent pandemics suggesting they are particularly important to human infection.

Previously covariant network analysis applied to other RNA and DNA viruses demonstrates observed covariance in viruses is often not an artifact of chance and likely influenced by imposed pressure, structure, and selection, including those provided by therapeutics (Aurora et al., 2009; Donlin et al., 2012; Sruthi and Prakash, 2019). Coevolving residues for large orthologous groups of proteins have also been useful in predicting structure (de Juan et al., 2013) and protein-protein interactions (Kamisetty et al., 2013) that could be targeted for drug interference. In the present study, we identified networks of co-varying amino acid residues in proteins encoded by nearly 13,600 CoV isolates. Because these interactions can be in theory critical to the CoV family of viruses based on their conservation in the evolution and adaption of these viruses to humans, we propose that they could define novel antigen and drug targets for CoV-related viruses. These targets are uniquely valuable because they are likely under strong selective pressure at two or more amino acid sites in one or more proteins; this property will make it less likely for a virus to mutant to escape immune responses or anti-viral drugs directed at protein sites enriched in such covariant-related targets. Finally, our data also provide evidence and a novel perspective not readily apparent in standard sequence alignment analysis for new adaptive recombination events and mutations that likely drove the emergence of SARS-CoV-related viruses into the human host. These insights could inform preparedness through therapeutic and vaccine development, as well as public health policy.

## Methods

Genome and protein sequences were downloaded from available NCBI and GISAID public databases. All genomes and accession numbers are provided in Supplemental Table S1. Protein sequences for genes in 1a,1b, S, 3a, E, M, and N were concatenated and aligned using CLC Bio Workbench (v8) for the 850 genomes using a gap open cost of 2.0 and a gap extension cost of 1.0 in very accurate mode and with MAFFT for the clinical SARS-CoV-2 strain using the default FFT-NS-2 setting (CLC-Bio Workbench 8, Nakamura et al., 2018). 13,611 clinical strains available on May 10, 2020 with significant contiguous coverage over the reference genes (>95%) were kept in the alignment. For Maximum Likelihood Phylogeny, we applied a WAG protein substitution model to our alignment and performed bootstrap analysis using 1000 replicates.

Pairwise and multiple residue covariance and scored were predicted using FastCov (Shen and Li, 2016). We set a purity score (0.96) for stringency cutoffs, but for clarity a raw table of predicted covariants is provided as Supplemental Table S2. This allowed binning of clusters and respective strains for Force Mapping in Gephi using the Multigravity ForceAtlas 2 setting and comparison of covariant residues based on clusters and strains (Jacomy et al., 2014). All clusters, strains, and residues are tabulated in Supplemental Table S3.

Clusters and residues were mapped in Gephi using the MultiGravity ForceAtlas 2 algorithm and this file is provided (Supplemental File F1) (Jacomy et al., 2014). The JavaScript Sigma.js library was used to export this as an interactive website.

Residues in S protein were mapped onto the PDB structure for the S trimer (6VXX.pdb and 6ACC.pdb) using PyMol (v.2.3.4) (PyMol, Song et al., 2018; Walls et al., 2020). Arpeggio was used to calculate interacting residues in the PDB file (Jubb et al., 2017). Circular graphing of key collections of residues was done using Circos (Krzywinski et al., 2009). Epitope mapping in the 3a NTD 44 amino acids was accomplished using BepiPred-2.0 (Jespersen et al., 2017)

## Acknowledgements

This work was supported by National Institutes of Health Grant AI-018045 to J.J.M. We would like to thank Matthew K. Waldor for his insight, perspective, and discussions about our analysis during the preparation of this manuscript.

## Supplemental Figures, Files, and Tables

**Supplemental Figure S1.**
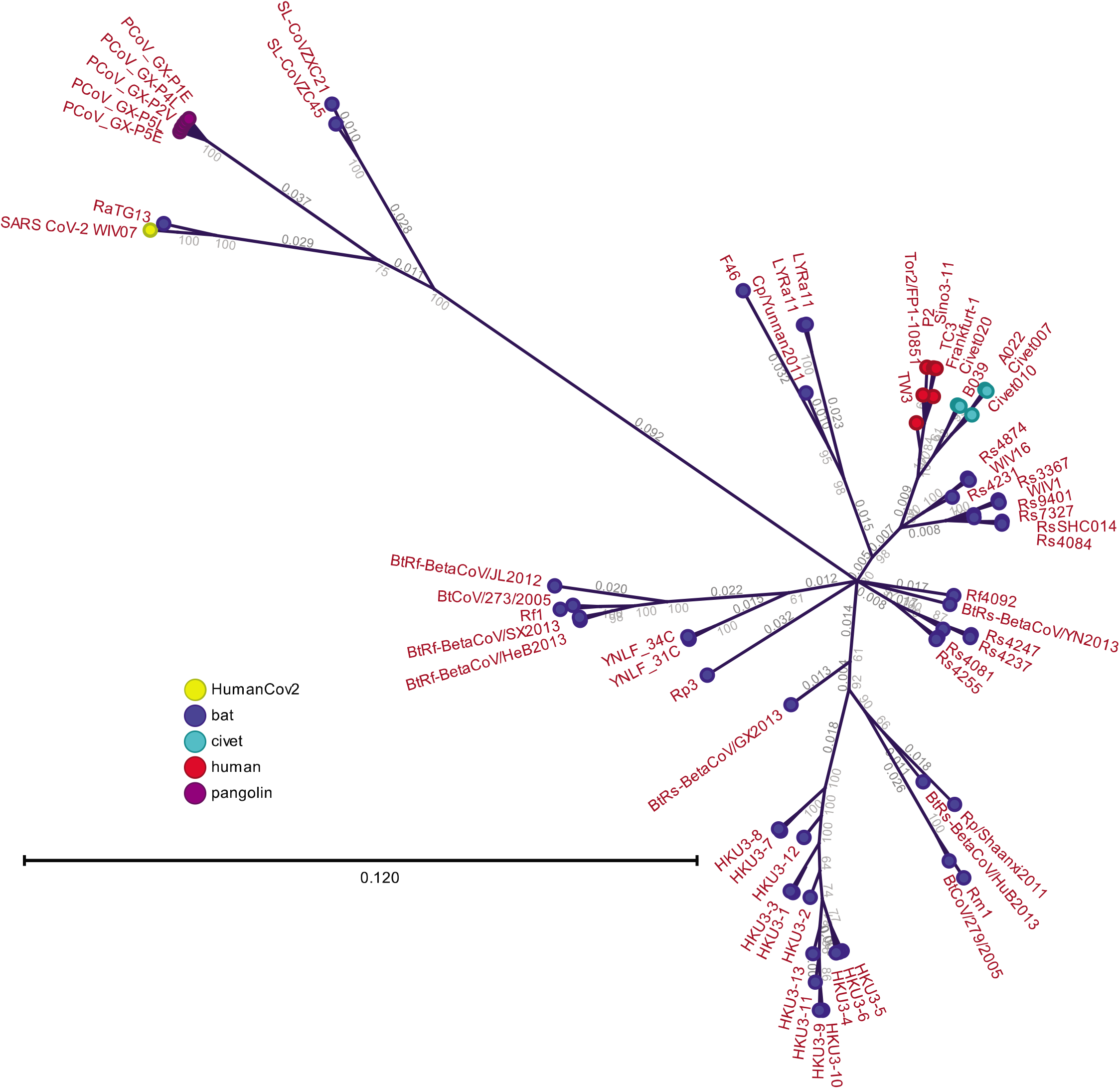
Phylogeny of the 850 CoVs. Maximum Likelihood Tree using the aligned amino acids from 1a/1b, S, 3a, M, E, and N proteins from bat, pangolin, civet, and human CoVs.

**Supplemental Figure S2.**
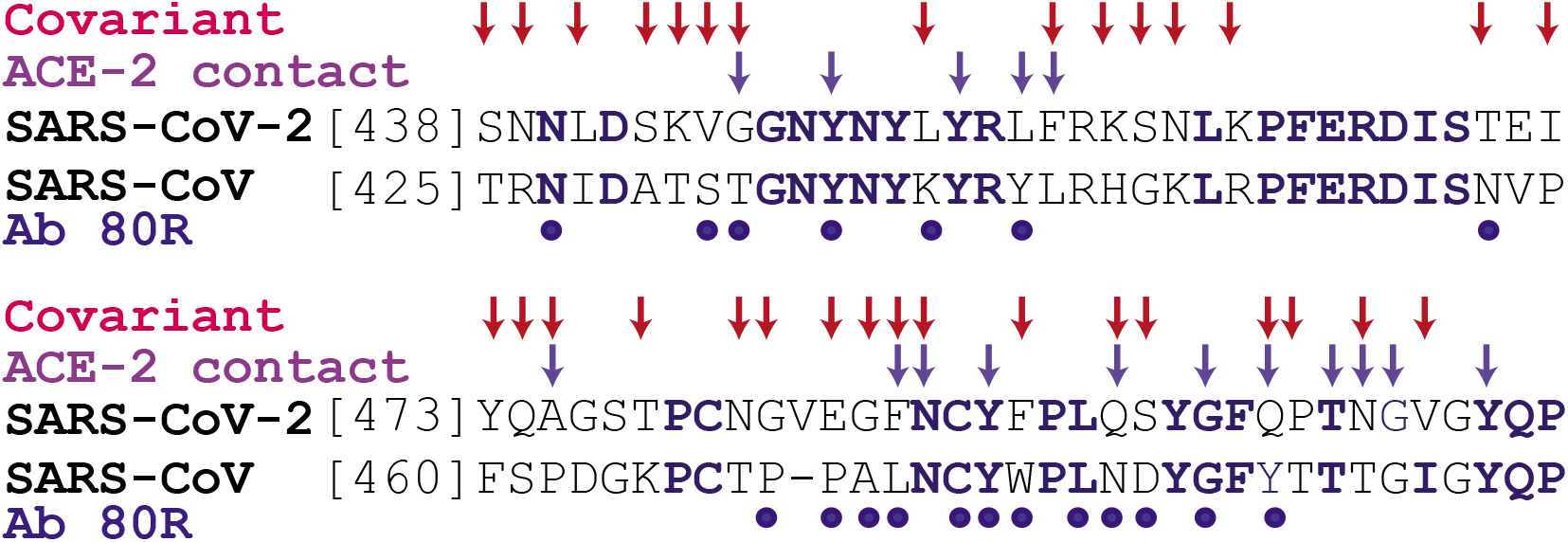
Alignment of covariants with the S receptor-binding domain. Aligned SAS-CoV and SARS-CoV-2 sequences in the S protein RBD showing sequence conservation (blue), covariant residues in SARS-CoV-2 (red arrows), determined contacts between both ACE-2 and Ab R80 and SARS-CoV S protein.

**Supplemental Figure S3.**
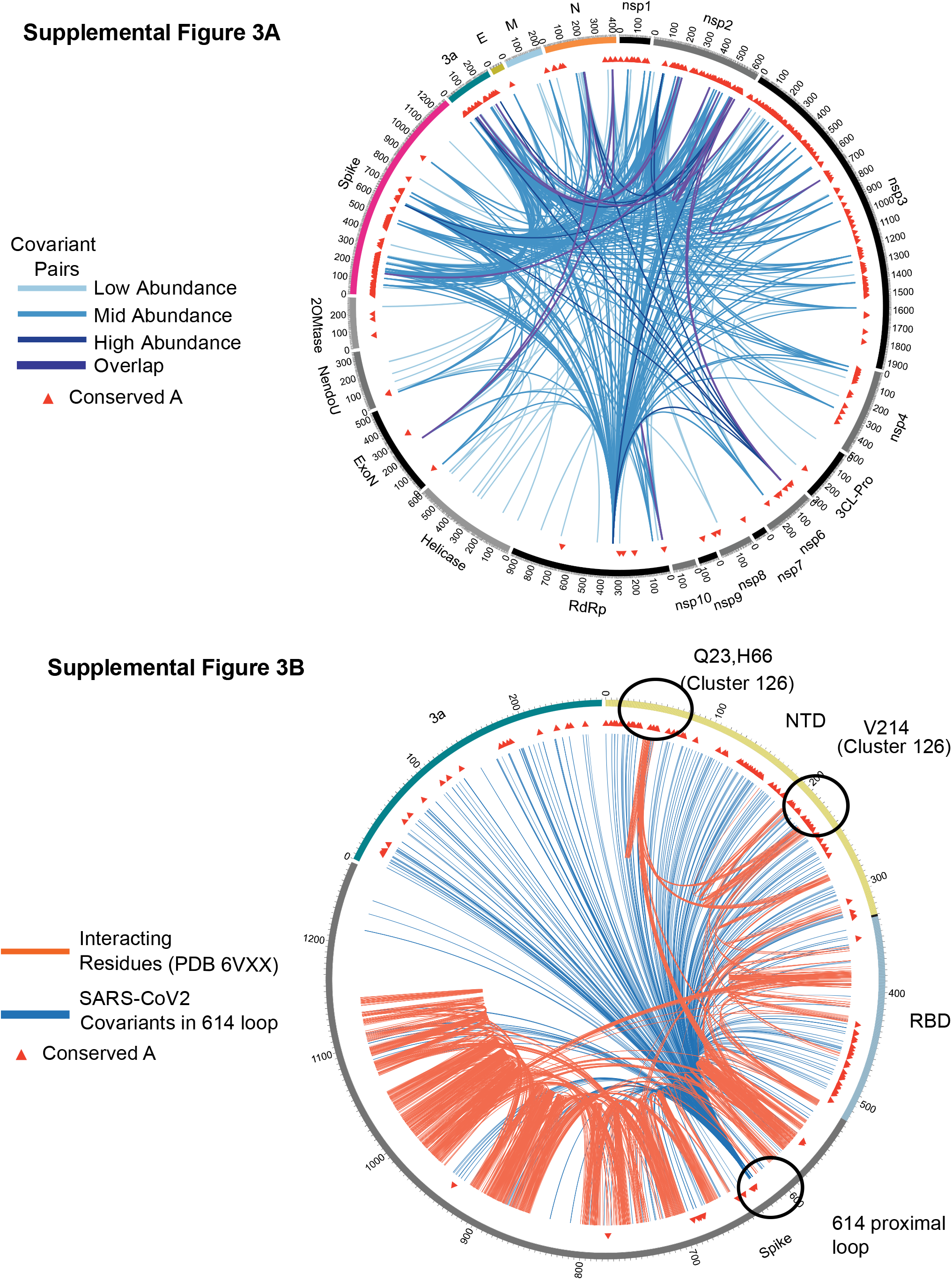
Plotting of predicted covariant and interacting residues in clinical strains and highlighted residues in the S protein. **(A)** 796 Covarying residue pairs detected in the a 13,611 clinical isolates. Abundance is indicated by hue. High abundance indicates both residues vary in at least 4% in these isolates, mid-abundance indicates one of these residue pairs is varied at least 4% in these isolates, and low-abundance is that neither are found in more than 4% of isolates. Overlap pairs are those covariant residues detected in both the 850 CoVs and the clinical strains. **(B)** Residues predicted to physically interact within the S protein from the PDB (6VXX) are orange. Residue linkages with S 614 and S and 3a proteins in the SAR-CoV-2 clusters are indicated in dark blue. The Location of residues identified in this work as emerging in both SARS-CoV and SARS-CoV-2 are circled in black.

**Supplemental Figure S4.**
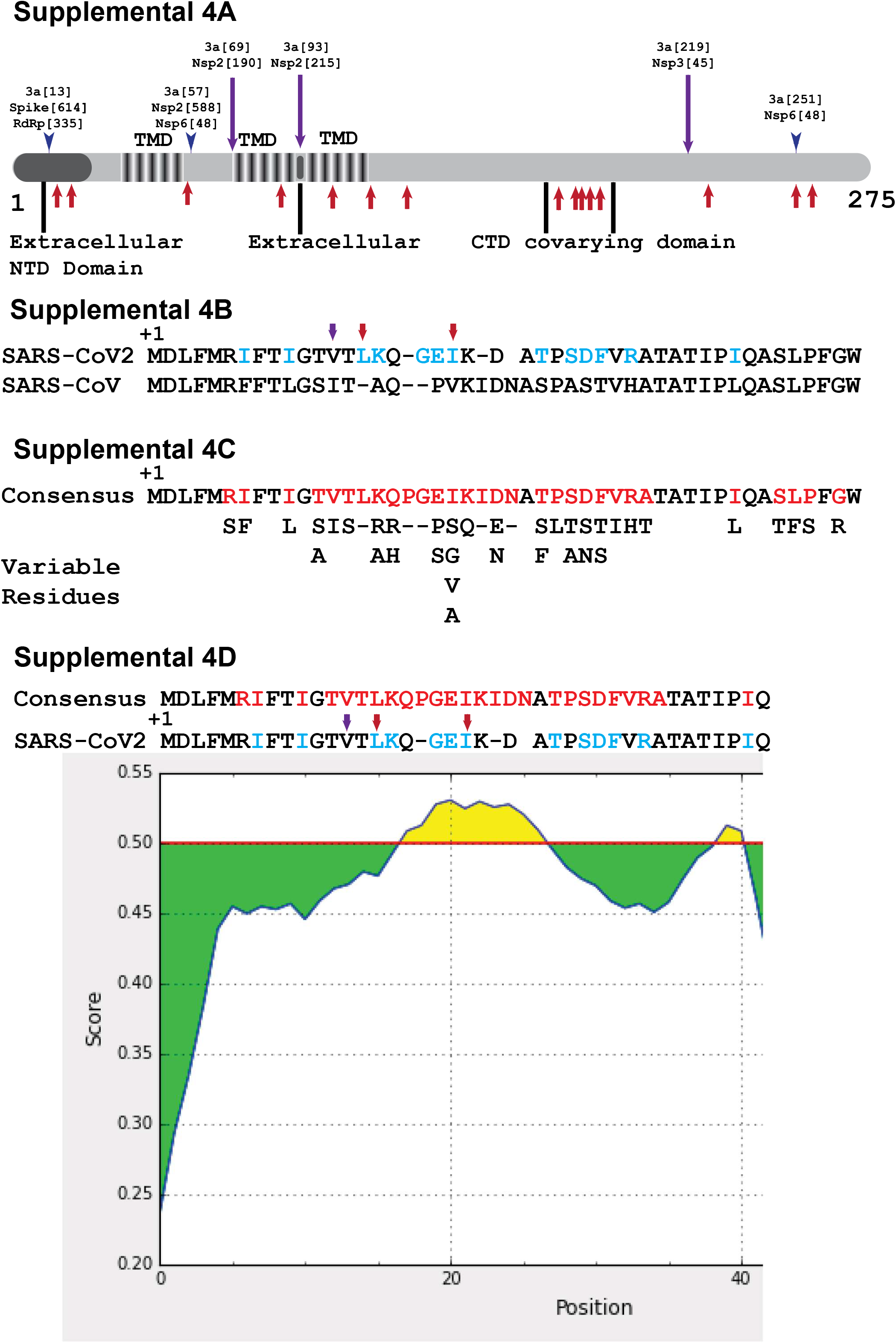
Covariant residues in 3a domains and observed variance in the 3a ectodomain. **(A)** Diagram showing the positions of extracellular, cytoplasmic, and transmembrane domains (TMD) in 3a protein. Red arrows indicate covariant residues from 850 CoVs and blue and purple arrows are those from clinical strains as shows in Supplemental S3. **(B)** Comparison of SARS-CoV-2 and SARS-CoV residues in the 3a N-terminal ectodomain and arrows from (A) showing covariance. Distinct residues are colored blue. **(C)** The consensus sequence of 3a protein derived from the alignment of CoVs in this work and substitutions observed each position. Variable residues are colored red. **(D)** BepiPred-2.0 epitope prediction using residues in the 3a ectodomain.

**Supplemental Figure S5.**
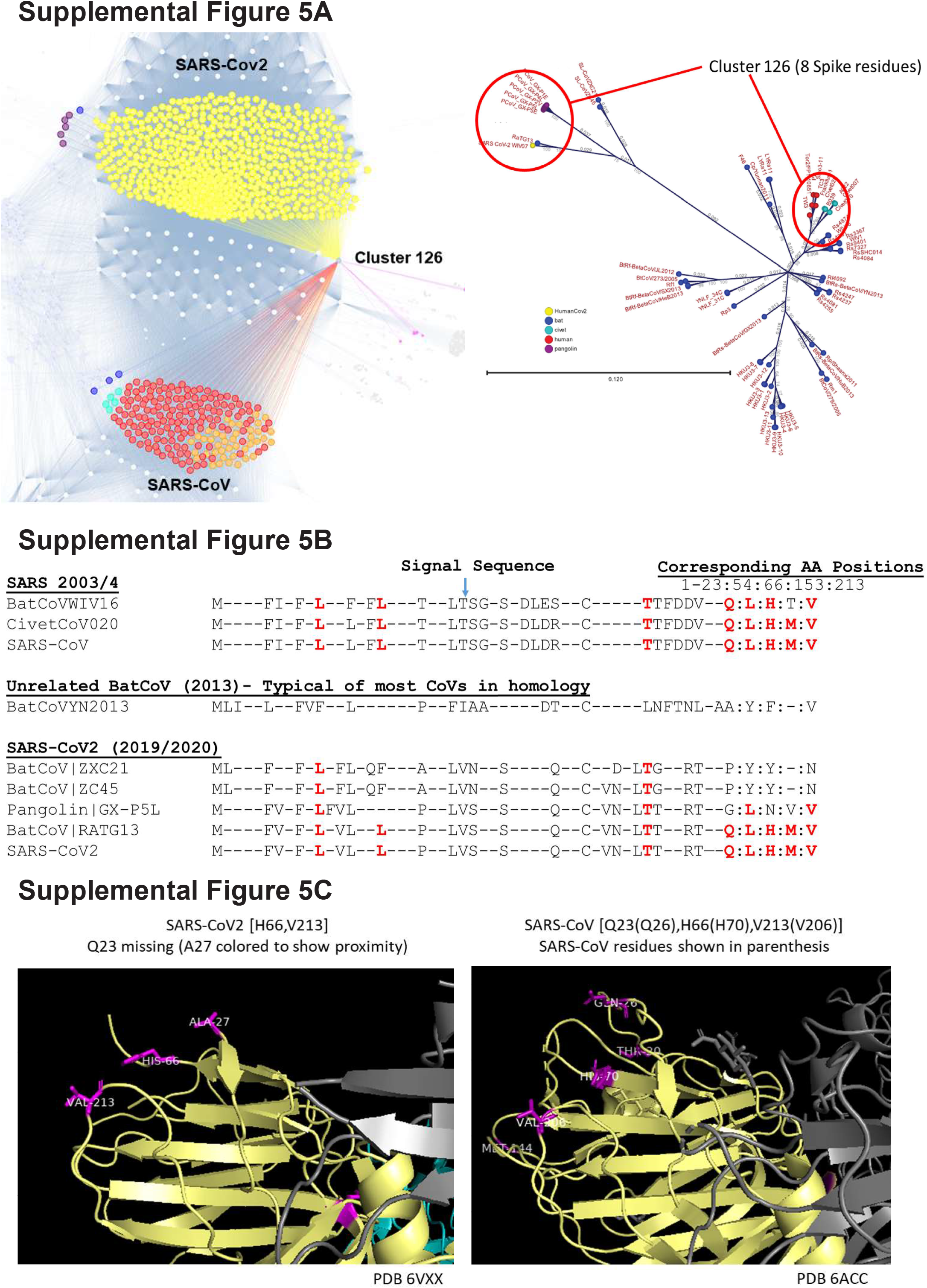
Covariants that emerged during SARS-CoV and SARS-CoV2. **(A)** Gephi Force map inset showing Cluster 126 and highlighted genomes that possess the Cluster 126 residues. The ML phylogenic tree showing the relative relationship of circled strains is included as a reference. **(B)** Alignment of strains to both SARS-CoV and SARS-CoV-2 showing the order and diversity of these 8 residues in S protein. **(C)** Structures of both SARS-CoV-2 (PDB 6VXX) and SARS-CoV-2 (6ACC) showing highlighted residues belonging to both. Q23 is missing from the SARS-CoV-2 and the closest residue A27 was labeled to provide an approximate location.

**Supplemental File F1. Gephi Force-Mapping File**

**Supplemental Table S1. Accession numbers and description of all genomes used in this work**

**Supplemental Table S2. Raw Covariance data and coordinates in both 850 CoV consensus and equivalent in SARS-CoV2. Amino acid positions are the best approximate based on alignment.**

**Supplemental Table S3. Cluster, Genome, and Residue linkage data used for force mapping.**

**Supplemental Table S4. Residue data by cluster used to chart Bat CoV and SARS-CoV-2 progression**

**Supplemental Table S5. 1231 Covariant residues in SARS-CoV-2, SARS-CoV Bat, Pangolin, and Civet.** Residues in the 850 CoV consensus and approximate positions in the SARS-CoV-2 proteins are indicated.

**Supplemental Table S6. Raw clinical pairwise covariance data and positions of residues in proteins.**

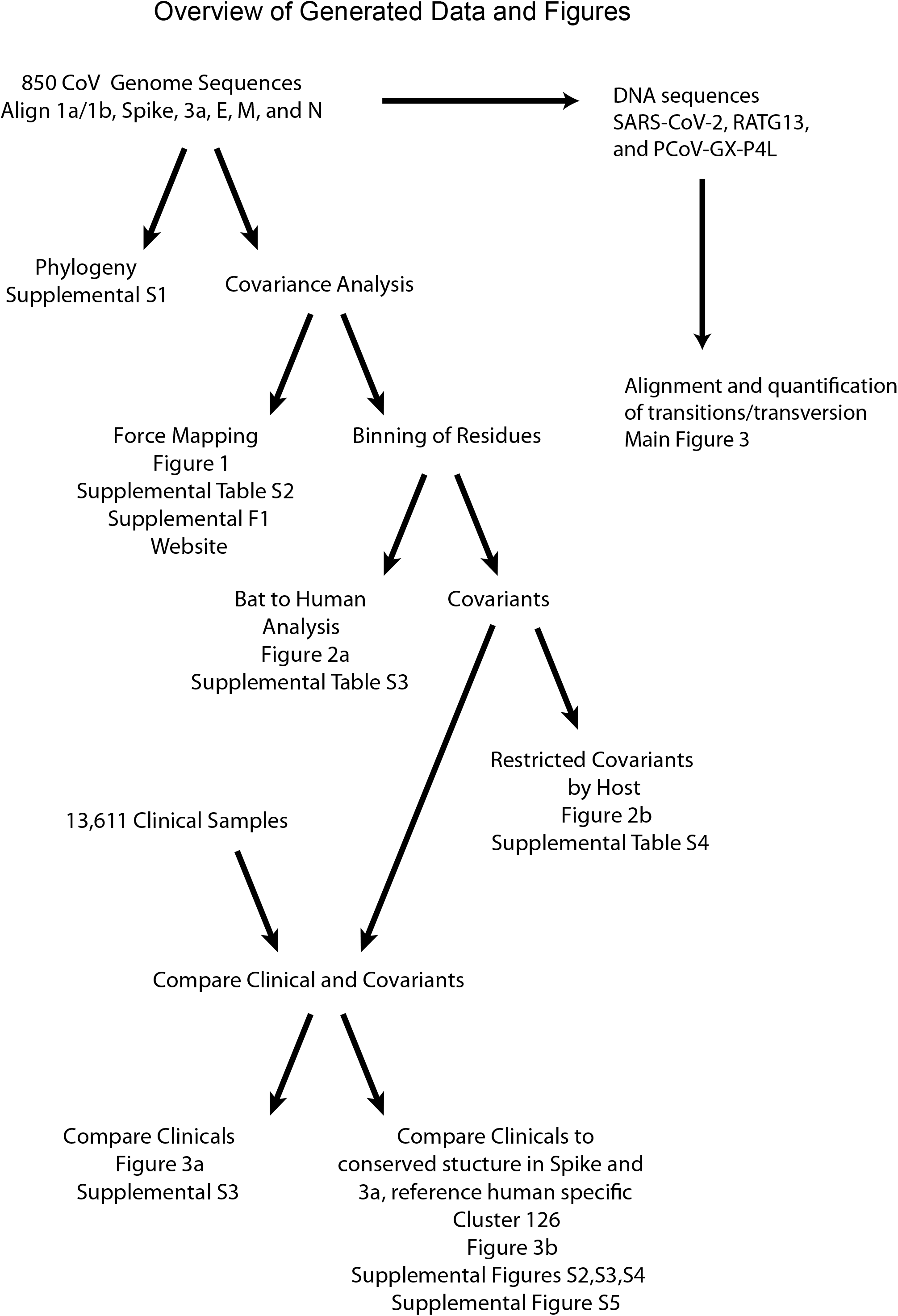

## Notes

### Competing Interest Statement

The authors have declared no competing interest.

